# Spatial neuronal synchronization and the waveform of oscillations: implications for EEG and MEG

**DOI:** 10.1101/401091

**Authors:** Natalie Schaworonkow, Vadim V. Nikulin

## Abstract

Neuronal oscillations are ubiquitous in the human brain and are implicated in virtually all brain functions. Often they are referred to by their frequency content, i.e., *α*-, *β*-, *γ*-oscillations. Although they indeed can be described by a prominent peak in the power spectrum, their waveform is not necessarily sinusoidal and shows a rather complex morphology which needs to be captured with multiple spectral harmonics. Both frequency and temporal descriptions of such non-sinusoidal neuronal oscillations can be utilized. However, in non-invasive EEG/MEG recordings the waveform of oscillations often takes a sinusoidal shape which in turn leads to a rather oversimplified view on oscillatory processes.

In this study, we show in simulations how spatial synchronization can mask non-sinusoidal features of the underlying rhythmic neuronal processes. Consequently, the degree of non-sinusoidality can serve as a measure of spatial synchronization. To confirm this empirically, we show that a mixture of EEG components is indeed associated with more sinusoidal oscillations compared to the waveform of oscillations in each constituent component. Using simulations, we also show that the spatial mixing of the non-sinusoidal neuronal signals strongly affects the amplitude ratio of the spectral harmonics constituting the waveform. This in turn has high relevance for the interpretation of the relative strength of spectral peaks, which is commonly used for inferring neuronal signatures corresponding to specific behavioral states.

Moreover, our simulations show how spatial mixing can affect the strength and even the direction of the amplitude coupling between constituent neuronal harmonics. Consistently with these simulations, we also demonstrate these effects in real EEG recordings. Our findings have far reaching implications for the neurophysiological interpretation of neuronal oscillations and cross-frequency interactions, as well as for the unequivocal determination of oscillatory phase.

## 1. Introduction

Neuronal oscillations are ubiquitous in the human brain, being present in both cortical and subcortical structures. Moreover, they have been shown to be relevant for sensory [1, 2], motor [3, 4] and cognitive [5, 6] functions. Traditionally, neuronal oscillations as recorded by EEG/MEG are considered to be sinusoidal. This observation is particularly driven by the analysis tools frequently used in neuroscience. These often include Fourier, Morlet wavelet and Gabor transforms, all of which use sinusoids as a basis function [7]. There is no a-priori reason why exactly these basis functions would be most relevant for describing neuronal oscillations. Many nonlinear periodic processes in nature are in fact quasi-sinusoidal [8]. For instance, the non-sinusoidal nature of ocean waves has for long time been recognized [9], where it was emphasized that conventional spectral analysis is not sensitive to the non-sinusoidal nature of periodic processes. Due to the complexity of such waves, analysis in time domain is often suggested and elaborate measures of horizontal and vertical asymmetries have been presented [10]. A similar claim has been recently voiced for large scale neuronal oscillations [11], which represent a particularly good example where many nonlinearities are present including thresholds, exponential decays and non-linear coupling between neuronal elements. It is therefore not surprising that often neuronal recordings only approximately resemble sinusoidal processes especially when they are obtained with invasive techniques [12, 13]. This in turn indicates that other concepts and analysis tools are needed for a more adequate description of periodic neuronal processes recorded with EEG/MEG.

Waveform was largely neglected in large scale EEG/ MEG analysis up until recently [14, 15]. However, the reasons why non-invasive neuronal recordings rather show sinusoidal oscillations in contrast to invasive recordings have not yet been clearly identified. Some evidence for non-sinusoidality is also visible in the spectral domain, as non-sinusoidal processes are manifested through the presence of additional peaks being usually integer multiples of the base frequency. Spectral harmonic peaks are often observed in LFP and EEG/MEG recordings. For instance, a spectral peak in *β*-frequency range has been found to be exactly twice the individual *α*-frequency peak [16, 17, 18].

The waveform of oscillations is also important for the understanding of non-linear neuronal interactions. which can be carried out not only within the same frequency band (e.g., *α*, *β*, *γ*) but also across different bands. In this case they are referred to as cross-frequency interactions and describe a mechanism through which spatially and spectrally distributed information can be integrated in the brain [19]. The extent to which the presence of such cross-frequency interactions can be due to spurious effects, particularly due to non-sinusoidal waveform of oscillations is being debated [20, 21, 22].

Furthermore, a description of oscillations which takes into account their non-sinusoidal waveform has implications for the understanding of oscillatory phase. Oscillatory phase is important in theories of neuronal processing [23, 24, 25], reflecting a change in membrane potential for many synchronous neurons. This in turn results in changes in cortical excitability, which has been associated with periodic inhibition. A non-sinusoidal waveform is associated with a deviation from a 50% duty cycle and with a non-uniform phase velocity [26]. This in turn would lead to non-uniform changes in cortical excitability and subthreshold stimulus detection rates along the oscillation cycle.

Here, we investigate measures for quantifying non-sinusoidality in the time domain, with simulation and analysis primarily focused on *α*- and *β*-oscillations in EEG recordings. The aim of the present study is to show that the degree of non-sinusoidality in oscillations may depend on the spatial mixing of the neuronal sources reflected in EEG/MEG/LFP recordings. Depending on synchronization strength and the temporal delay between neuronal populations, the resulting waveform of oscillations can vary from strongly non-sinusoidal to sinusoidal. Spatial mixing will influence measures such as amplitude envelope correlations and *α*/*β*-ratio, as different temporal delays will cancel or enhance different frequency components of the non-sinusoidal waveform. Moreover, this might lead to spurious inferences about cross-frequency interactions, which may rather relate to changes in the waveform reflecting in turn changes in spatial synchronization.

## 2. Materials and Methods

### 2.1. Experimental Recordings

#### 2.1.1. Participants

The study protocol conformed to the Declaration of Helsinki and by the ethics committee at the medical faculty of the University of Leipzig (reference number 154/13-ff). The EEG data were previously collected as part of the “Leipzig Cohort for Mind-Body-Emotion Interactions” data set (LEMON) which is currently being prepared for release. Written informed consent was obtained prior to the experiment from all participants. Data from 13 participants were excluded due to missing event information, different sampling rate, mismatching header files or insufficient data quality. Additionally, data from 17 participants was excluded for insufficient signal-to-noise ratio (see section Data analysis and Statistics). This resulted in data sets from 186 participants (117 male, 69 female, age range: 20–70 years) with no history of neurological disease and usage of CNS drugs.

#### 2.1.2. EEG setup

Scalp EEG was recorded from a 62-channel active electrode cap (ActiCAP, Brain Products GmbH, Germany), with 61 channels in the international 10-20 system arrangement and one additional electrode below the right eye recording vertical eye movements. The reference electrode was located at electrode position FCz, the ground was located at the sternum. Electrode impedance was kept below 5 kΩ. Data were acquired with a BrainAmp MR plus amplifier (Brain Products GmbH, Germany) at an amplitude resolution of 0.1 *μ*V with a bandpass filter between 0.015 Hz and 1 kHz and with a sample rate of 2500 Hz. The recordings were performed in a sound attenuated EEG booth.

The experimental session was divided into 16 blocks, each lasting 60 s, with two conditions interleaved, eyes closed (EC) and eyes open (EO), starting in the EC condition. Participants were instructed to fixate on a digital fixation cross during EO blocks. Changes between blocks were announced with the software Presentation (v16.5, Neurobehavioral Systems Inc., USA). Only data from the EC condition were used for analysis.

### 2.2. Data analysis and computational modelling

#### 2.2.1. Measures for assessing non-sinusoidality

To exploit the vast richness of the momentary EEG-signal, we utilize measures of waveform shape in the raw signal with only limited band-pass filtering. The waveform features of an asymmetric signal are illustrated in Fig. 1. The crest period T_c_ is defined as the time from up-crossing to the next down-crossing. Conversely, the trough period T_t_ is the time from down-crossing to next-up-crossing. Each period is associated with two amplitude values, the crest amplitude A_c_ and the trough amplitude A_t_. We propose to assess the non-sinusoidality of a signal by considering the ratio of the crest period versus the trough period 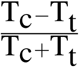, termed CT-difference. The more this value deviates from 0, the more non-sinusoidal the signal is. For more stable estimation, this can be done over several segments of data, with 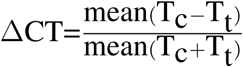. To compute T_c_ and T_t_-values for empirical as well as synthetic data, we used the WAFO toolbox [10], originally developed for the analysis of ocean waves. For computation of ΔCT, EEG data were bandpass filtered in the frequency band 3–45 Hz (Butterworth, filter order = 4). We used all periods with associated pooled crest and trough amplitudes larger than the 50^th^ amplitude percentile in order to avoid a contamination with 1/f-noise in EEG signals. Similar measures for the description of oscillatory waveform have also been proposed by Cole and Voytek [14].

**Figure 1:**
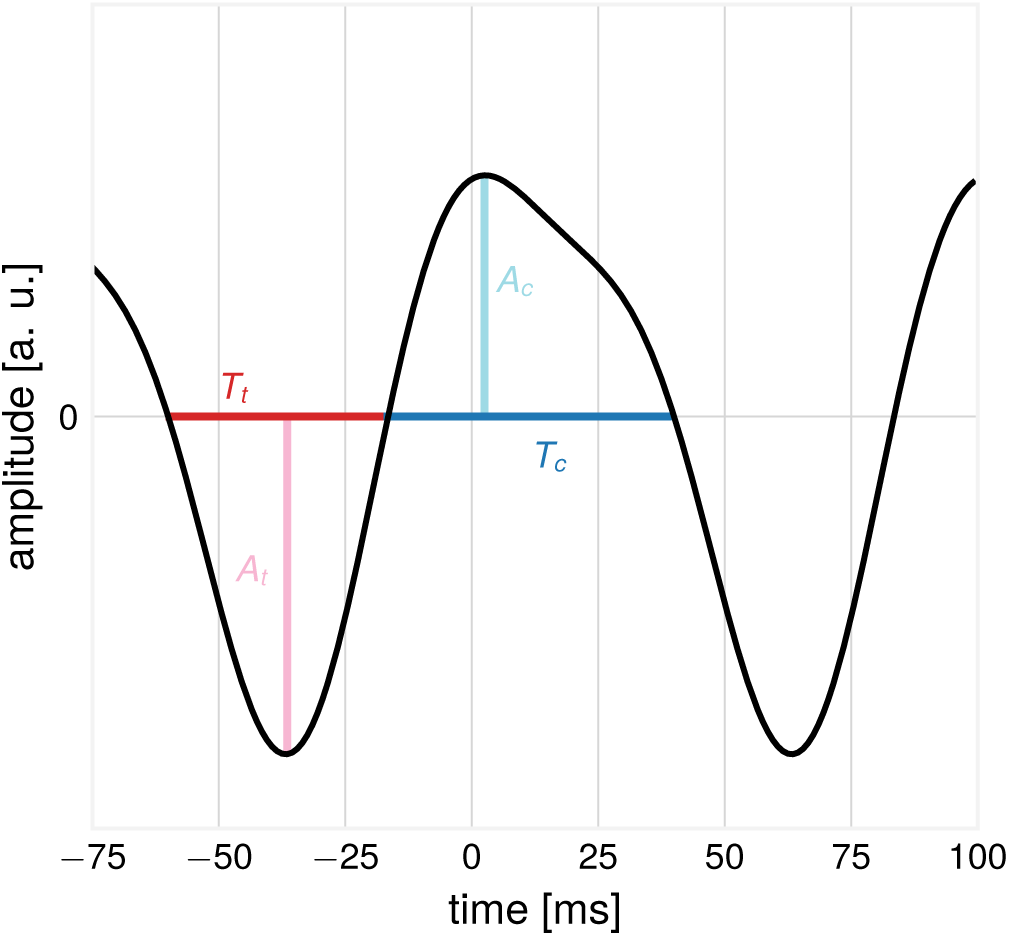
Illustration: features of a non-sinusoidal waveform. The trough period T_t_ and the associated trough amplitude A_t_ and the crest period T_c_ and associated crest amplitude A_c_.

#### 2.2.2. Time series simulations

To study properties of the proposed measures, we employed the following simulation procedure: First, basis functions with an arc-shape waveform were constructed, resembling the non-sinusoidal activity of a source. The waveform is composed of two sinusoids, the *α*-component with the base frequency of 10 Hz, and the *β*-component with a frequency of 20 Hz [27]. The sinusoids have a fixed phase shift relative to each other, the power of the *β*-component is four times smaller than the *α*-component: *μ*(*t*) = *A*_1_ · sin(*f* · 2*π · t*) + *A*_2_ · sin(2*f* · 2*π · t* + *ψ*), with *A*_1_ = 1*, A*_2_ = 0.25*, f* = 10 Hz*, ψ* = 1. The waveform is asymmetric by construction, see Fig. 1 for the waveform and corresponding features. In a second step, signals from *N* sources which have a temporal shift *ϕ*_*i*_ relative to each other, with *ϕ*_*i*_ ∼𝒩 (0*, σ*) were added to result in a compound signal 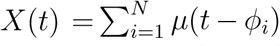. The sources represent spatially close neuronal populations participating in the generation of the compound signal. In some cases, the basis signals were amplitude-modulated (resulting in the same amount of amplitude modulation for *α* and *β*-components) to produce amplitude envelopes with a 1/f-distribution as found in real EEG recordings.

### 2.3. Simulation of α/β-ratios and amplitude envelopes

*α*/*β*-ratios and amplitude envelope correlations were evaluated using the compound signal *X*(*t*). The compound signal was composed of 20 sources, mixed with temporal delays drawn from a normal distribution with mean 0 and varying values for the standard deviation *σ*. *α*/*β*-ratios were calculated as the ratios of *α*- and *β*-SNR values of the compound signal, evaluated over time series segments of varying length. Here, *α*-SNR was taken as oscillatory power at the *α*-peak frequency and *β*-SNR as the oscillatory power at twice that frequency. Power was computed by FFT, Hann window, 1 s window length, 50% overlap. To investigate time courses between *α*- and *β*-components of the compound signal, we calculate correlations between their amplitude envelopes. Amplitude envelopes were calculated by individually bandpass-filtering the compound signal in the base frequency range and first harmonic frequency range *±* 2 Hz, respectively (Butterworth, filter order = 9). Amplitudes envelopes were determined for each frequency band by the means of the Hilbert transform. Then, the Spearman rank correlation coefficient was calculated between *α*- and *β*-amplitude envelopes. The calculation was repeated 1000 times, every time using a new instantiation of the compound signal, sampling new temporal shifts and 1/f-noise for amplitude modulation.

#### 2.3.1. EEG data analysis and Statistics

The BBCI toolbox [28] was used for EEG data analysis. The data were downsampled from 2500 Hz to 250 Hz, bandpass filtered in the frequency range 1–45 Hz (Butterworth filter, filter order 4). Visual inspection was utilized to exclude outlier channels with frequency shifts in voltage and poor signal quality and data intervals with extreme peak-to-peak deflections or large bursts of high frequency activity. Principal component analysis (PCA) was used for dimensionality reduction by keeping PCs that explain 95% of the total data variance. Next, independent component analysis (ICA) based on the Infomax algorithm was performed. Components reflecting eye movement, eye blink or heartbeat related artifacts were removed. Remaining independent components (mean number: 21.4, range: 14–28) were projected back to sensor space for further analysis.

As we are interested in oscillatory activity, only participants with sufficient signal-to-noise ratio in the *α*-band were included. To determine this, EEG time series were spatially filtered with a Laplacian filter, and the frequency spectrum (FFT, Hann window, 1 s window length, 50% overlap) was computed. The SNR-values of spectral peaks in the *α*-band (8–13 Hz) were considered with the 1/f-component removed by fitting a polynomial function to the computed spectrum and subtracting the estimated 1/f-fit [27]. Participants were included if at least one channel displayed a SNR > 5 dB in the *α*-band, as evaluated over the whole recording length.

The LEMON data set was available with sampling frequency of 250 Hz. To improve estimation of zerocrossing timing, the data was interpolated to a sampling frequency of 1000 Hz (spline interpolation), for 1 millisecond precision of T_c_ and T_t_-values. For the extraction of oscillatory components, spatial-spectral decomposition (SSD) [29] was used. The frequency band of interest was identified as the subject-individual spectral peak in *α*-frequency range *±* 2 Hz. All SSD components with SNR > 5 dB were kept. For demonstrations, we generated synthetic compound signals from data by adding extracted SSD components with a varying time shift. Empirical *α*/*β*-ratios were calculated as the ratio of *α*- and *β*-SNR values, evaluated over segments of varying time length. The subtraction of 1/f-fit was not performed here, as its estimation becomes unstable for segments of short length. Amplitude envelopes were calculated with the same parameters as for the synthetic compound signals.

## 3. Results

### 3.1. Waveforms become more sinusoidal with decreased spatial synchronization

Spatial mixing of non-sinusoidal sources results in more sinusoidal compound signals. Considering the example in Fig. 2a, seven basis signals are added with temporal delays drawn from a normal distribution. The compound mean signal has lost its asymmetrical shape and shows no difference between crest and trough periods (shown for one oscillation cycle in Fig. 2b), compared to the basis functions. Note that the disappearance of the non-sinusoidal waveform is not due to the changes in SNR but due to the time delay between individual sources. As a temporal delay of e.g. 10 ms is equivalent to 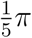 for the *α*-component, but twice as large, 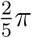 for the *β*-component, this leads to faster attenuation of the *β*-component.

**Figure 2:**
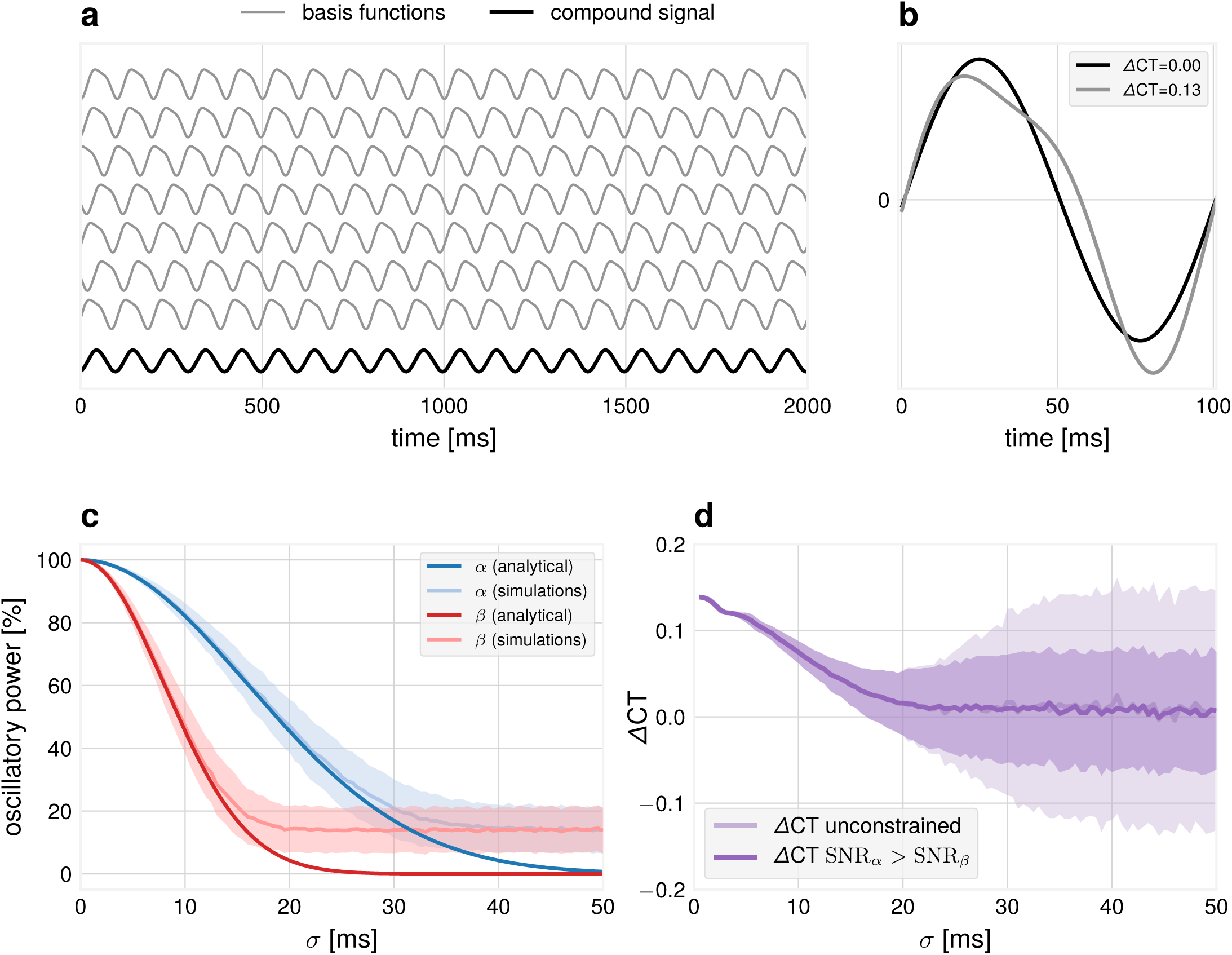
Simulation: dependence of waveform measures on spatial mixing. (a) Illustration how non-sinusoidal waveforms add up to a more sinusoidal compound signal if they are shifted with respect to each other with a certain standard deviation *σ* (example for *σ* = 30 ms). (b) Examples of one cycle of source and compound waveforms and their respective zero-crossings with associated ΔCT values. (c) The relative power of the two frequency components of the compound signal as evaluated from the Fourier spectrum. The *β*-component attenuates faster than the *α*-component. Number of iterations = 1000. Error bars indicate *±* 1 SD. In simulations, the obtained power for large temporal delays is constrained by the finite number of generators used and results in a deviation from the analytical solution. (d) ΔCT for the compound signal drops as a function of the standard deviation *σ*. In unconstrained simulations, SNR_*β*_ can get larger than SNR_*α*_, resulting in high values of ΔCT. Number of iterations = 1000. Error bars indicate *±* 1 SD.

To quantify the attenuation of the faster component, we computed the power spectrum of the compound signal *X*(*t*) by the Fourier transform as a function of standard deviation of the temporal delays *σ*. The analytical solution is proportional to exp(2 (*π · σ · f · N*)^2^), as obtained by Fourier analysis of the compound signal as a function of *σ*. The quadratic dependency on the frequency term results in a faster attenuation of higher frequencies, as seen in Fig. 2c. This results in a more sinusoidal signal for larger values of *σ*. Not only spectral power, but also the proposed measure for non-sinusoidality in the temporal domain is able to detect non-sinusoidality in the compound signal as a deviation from 0 (see Fig. 2d). In our unconstrained simulations, the spectral peak of the *β*-component can be higher than the spectral peak of the *α*-component (see also later sections in the results). In this case, extreme ΔCT values are observed, leading to an increased standard deviation for large temporal delays. The implication is that the degree of non-sinusoidality present in the waveform can serve as an indicator of spatial synchrony. It can also constrain the mixing coefficients, which are known in simulations, but are not known for real EEG recordings.

As an example for the ΔCT-measure, we illustrate the T_c_ and T_t_-distributions for different types of EEG oscillations in the 8–13 Hz frequency band for an exemplary participant. After SSD decomposition, one motor and one visual component was identified from the associated activation patterns. We compute T_c_ and T_t_ for motor and posterior oscillations, shown in Fig. 3. In this participant, a more non-sinusoidal oscillation can be found for the motor-component.

**Figure 3:**
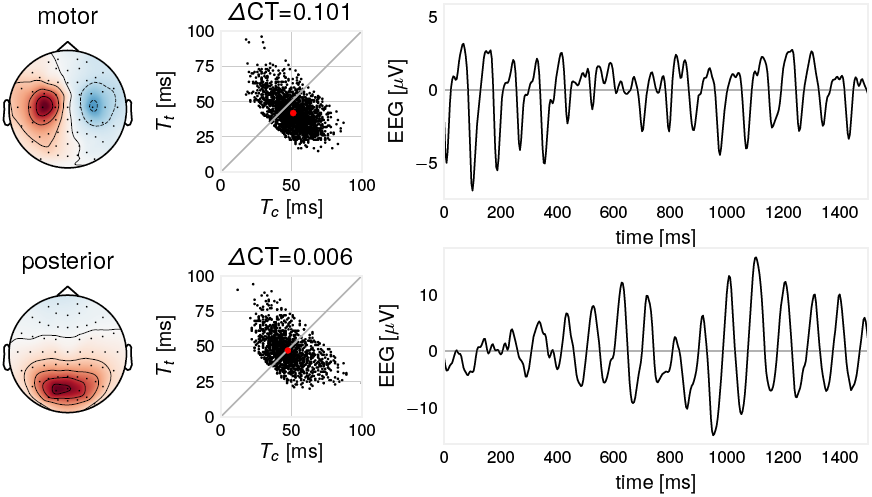
Illustration: ΔCT differs for two EEG oscillatory components. Components extracted for one participant. Left: SSD component pattern. Middle: for every oscillatory cycle, there are two corresponding ΔCT and T_c_and T_t_-values. Red dot indicates mean T_c_and T_t_-values. Right: example time course excerpt of the signal. In this case, the motor component shows a characteristic arc-like shape with larger non-sinusoidality than the posterior component.

### 3.2. Demixed recordings show higher degree of non-sinusoidality

We quantified the extent to which ΔCT is affected by a demixing procedure, which brings sensor signals closer to their sources. For this, ΔCT was computed in sensor space recordings for all included participants, as well as for SSD-extracted components. SSD components have a higher ΔCT indicating a higher degree of non-sinusoidality across participants (p=5.7 · 10^−15^, two-sided Wilcoxon signed rank test), as illustrated in Fig. 4.

**Figure 4:**
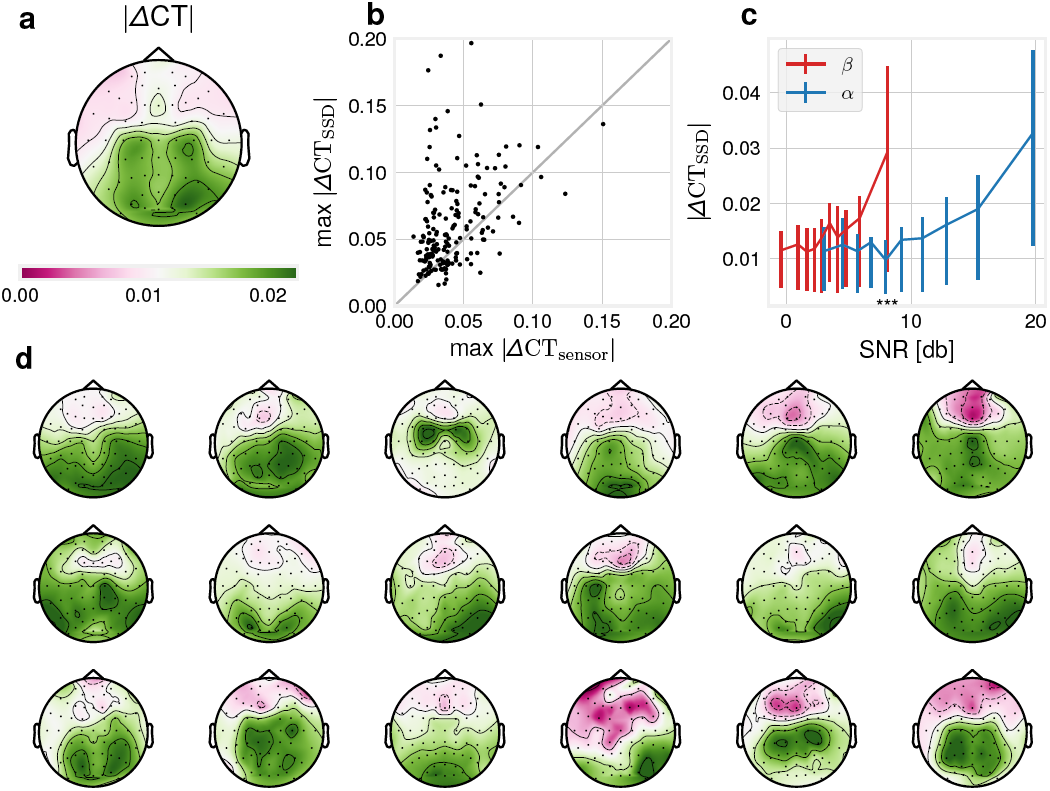
Data analysis: empirical ΔCT-distributions. (a) absolute ΔCT across participants computed from sensor space data, plotted topographically. (b) Maximal SSD absolute ΔCT is larger than sensor space absolute ΔCT across participants, N=186, p=5.7 10^−15^, two-sided Wilcoxon signed rank test. (c) Binned *α*-SNR and *β*-SNR as estimated from 1/f-adjusted spectrum versus mean absolute ΔCT in that bin. Error bars are 25^th^ − 75^th^ percentile value ranges for absolute ΔCT for the respective bin. Wilcoxon signed rank test between absolute ΔCT-values corresponding to the 10^th^ bin of *β*-SNR and 5^th^ bin of *α*-SNR: p-values: 5.40 *·* 10^−18^. (d) 18 single subject absolute ΔCT-topoplot examples show substantial variability of spatial ΔCT-distribution. Participants were selected according to the number of channels satisfying the SNR-criterion of 5 dB, so a topography is visible.

The dependence of the ΔCT of SSD-components to SNR was assessed by computing the SNR in the *α*-frequency band via 1/f-corrected spectrum and absolute value of ΔCT for all SSD-components with *α*-SNR > 5 dB. We found a correlation of .242 (Spearman’s rho, *p* < 6.99 *·* 10^−31^) of absolute ΔCT with *α*-SNR, with more non-sinusoidal signals as measured by ΔCT for higher SNR. Resorting the absolute ΔCT-values according to their associated *β*-SNR values shows that a *β*-SNR-level of the same magnitude as *α*-SNR of e.g ∼ 8 dB is associated with higher ΔCT values. In other terms, a pronounced *β*-peak in the 1/f-corrected spectrum corresponds to a higher degree of non-sinusoidality than an *α*-peak of the same magnitude. This observation is in agreement with our simulations presented above indicating that the presence of *β*-oscillations defines non-sinusoidality of the waveform.

The topographic distribution of ΔCT-values can be seen in Fig. 4d, which shows considerable variation across participants. Although the group average in Fig. 4a shows increased values for both central-motor and occipital channels, on a single subject level either a central-motor or an occipital maximum is rather visible. In sum, the non-sinusoidality of EEG recordings is affected by spatial mixing of oscillatory sources and also by SNR in relation to 1/f-noise.

### 3.3. Constructive and destructive interference with respect to temporal delays

A spatial summation of basis signals with the same spectral content but different temporal delays can have differential consequences for the respective constituent frequencies, enhancing or diminishing respective oscillatory power. We provide three examples for this phenomenon.

#### 3.3.1. Attenuation of the α-component and enhancement of the β-component

Non-sinusoidal signals can mix in such a way that the more prominent *α*-component is attenuated, while higher harmonics are preserved. This will lead to the emergence of *β*-events in the compound signal without the strong presence of *α*-events, even though the original source signals still have a high amount of *α*-frequency spectral content. Fig. 5 shows two basis segments of real EEG recordings which have the same spectral content (the bottom one is the time-reversed version of the top one), which are added with varying temporal delays. Depending on this delay, this results in periods in the compound signal where the *α*-component is diminished, and a higher amount of *β*-spectral content emerges. This phenomenon is most pronounced if the temporal delay is approaching *π* of the base oscillation (i.e. destructive interference), which corresponds to 2*π* for the first harmonic (i.e. constructive interference)

**Figure 5:**
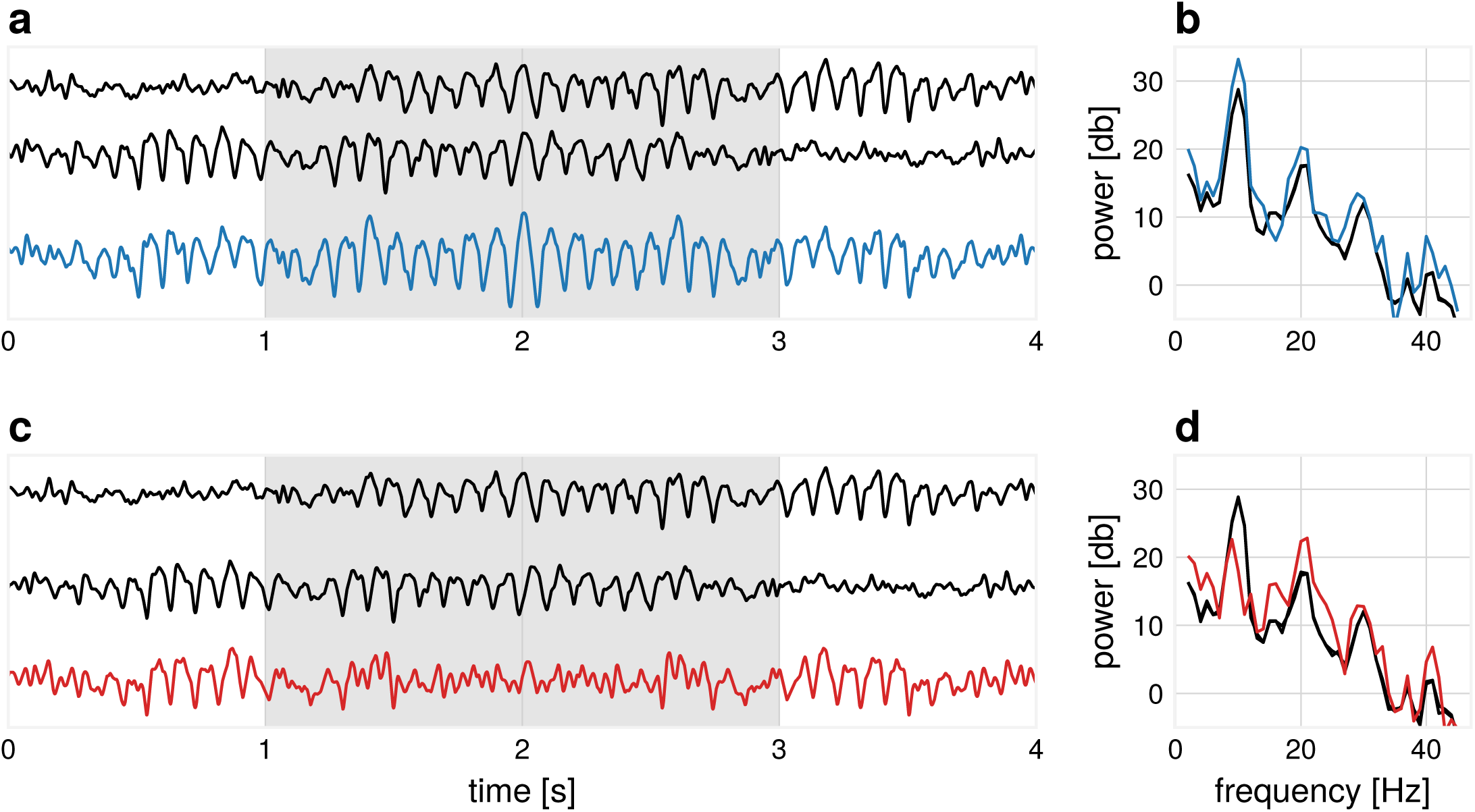
Illustration: the emergence of *β*-events from *α*-dominated sources. (a) Two basis functions (black) mix with a temporal delay (*σ* = 15 ms) such that the *α*-power is enhanced in the compound signal (blue) during the segment marked in gray. (b) The corresponding power spectrum of the segment marked in gray for subplot (a) for basis functions (black) and compound signal, where both *α*- and *β*-spectral peaks are enhanced (blue). (c) Two basis functions (black) mix with a temporal delay (*σ* = 43 ms) such that the *α*-power is diminished in the compound signal (red) during the segment marked in gray. (d) The corresponding power spectrum of the segment marked in gray for subplot (c) for basis functions (black) and compound signal, where the *α*-peak is largely diminished and *β*-peak is enhanced (red).

#### 3.3.2. Influence of temporal delays on α/β-ratios

Next, we show the impact of spatial synchronization on *α*/*β*-ratios in a simulation with a higher number of source basis signals, each having identical spectral content. As the temporal delay between source signals increases, a spread in *α*- and *β*-SNR becomes visible, as shown in Fig. 6.

**Figure 6:**
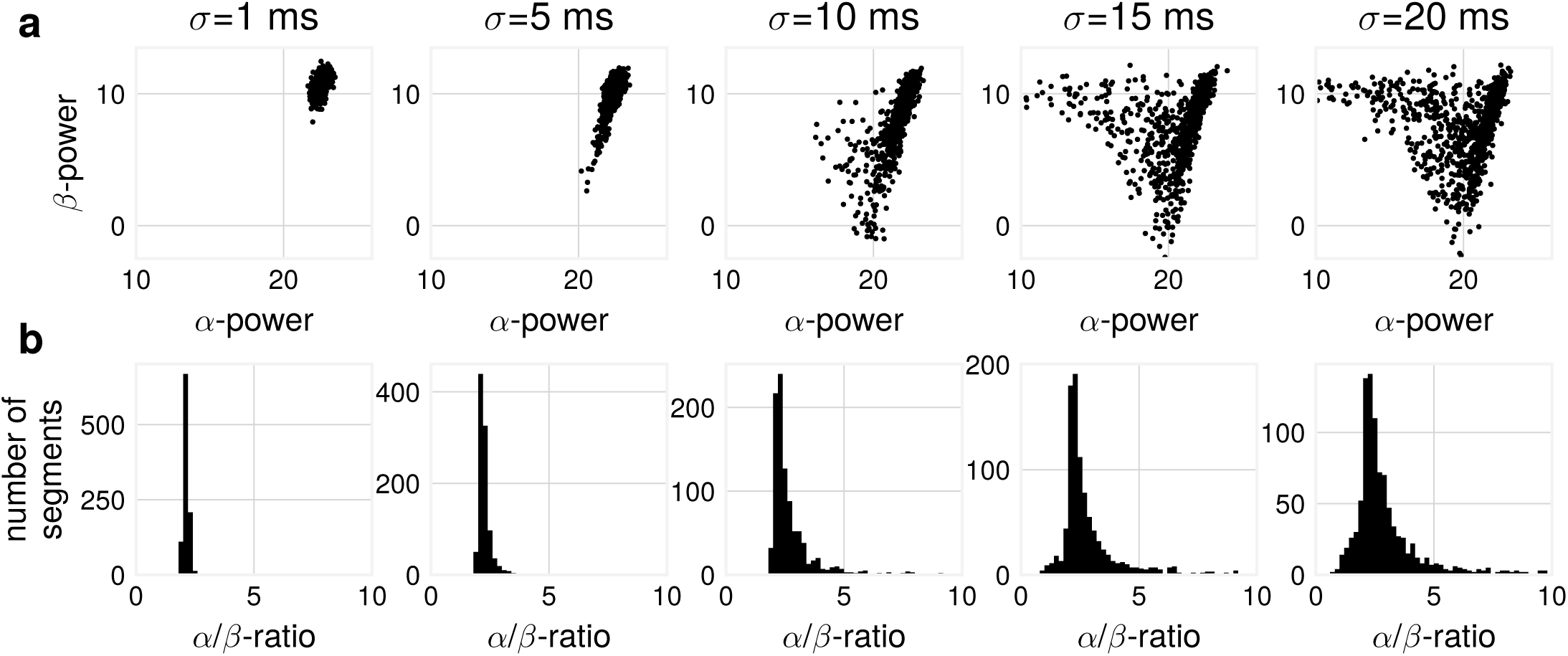
Simulation: *α*/*β*-ratios change with different levels of spatial synchronization. (a) *α*- vs. *β*-power for compound signals composed of non-sinusoidal basis signals for varying standard deviations of the time shift. Spread in power is larger for larger time shifts. Each point signifies *α*- and *β*-power as computed from one compound signal. (b) This results also in a spread of *α*/*β*-ratios as constructed from *α*- and *β*-power. Number of generators: 20, number of iterations: 1000, segment length = 5 s.

A mixture of a larger number of source signals can yield a change in spectral content of the compound signal, even without any changes in the spectral content of the source basis signals. This has implications for measures relating oscillatory power of two frequencies, for instance *α*/*β*-ratio. Changes in these measures may not necessarily reflect changes in spectral content, but a change in temporal coupling of non-sinusoidal signals.

*α*/*β*-ratios were also computed for real EEG recordings. A spread in these ratios is visible, see Fig. 7a. Considering segments of short length, periods were *β*-power is larger than *α*-power results in *α*/*β*-ratios < 1. To summarize, in agreement with the simulations, we show that obtaining larger *β*-power than *α*-power is also possible in real EEG data and this phenomenon can be observed more often when considering shorter segments (Fig. 7b).

**Figure 7:**
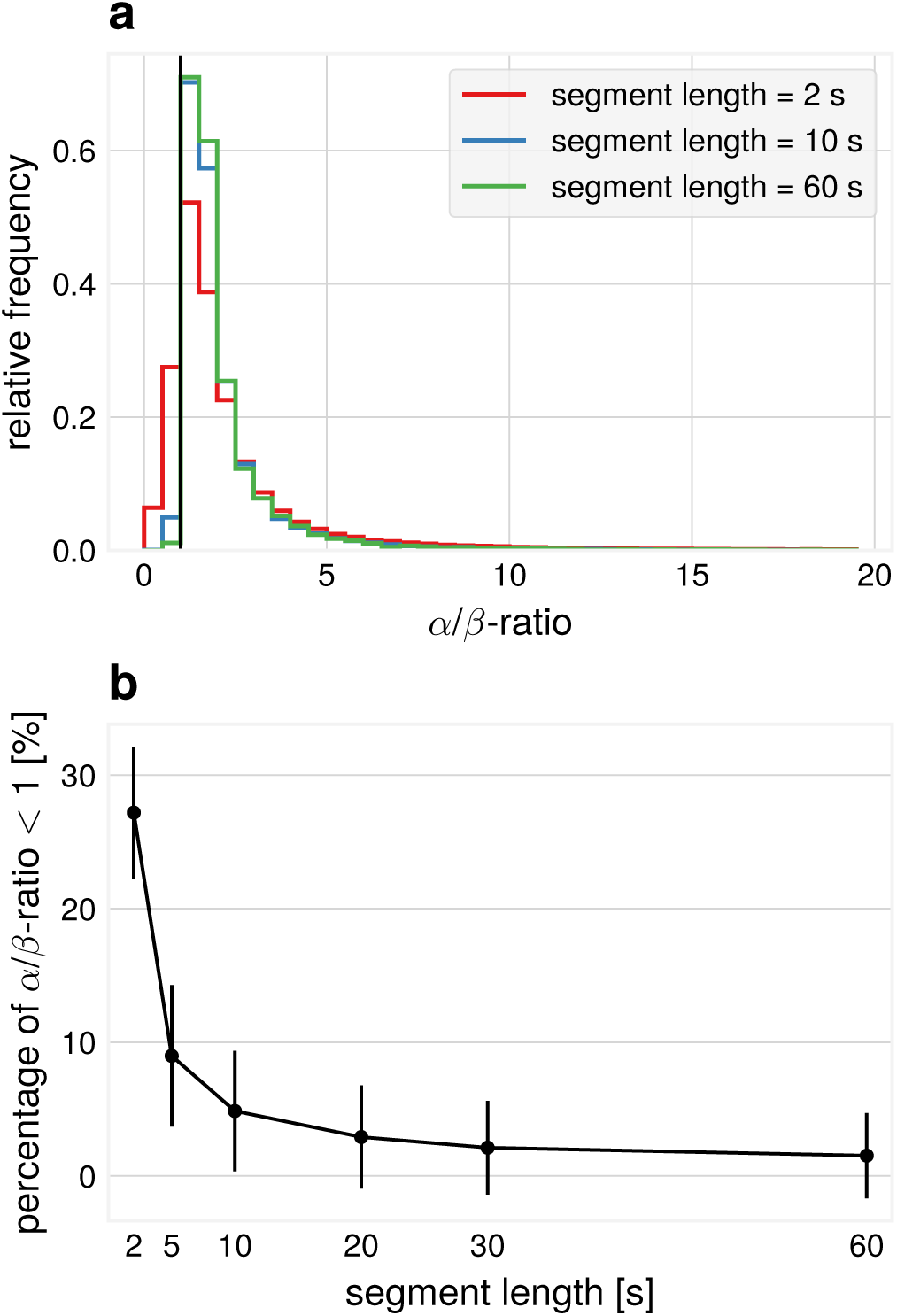
Data analysis: empirical *α*/*β*-ratios. All SSD components from all participants pooled. (a) *α*/*β*-ratios for different segment lengths, pooled over segments. (b) Percentage of *α*/*β*-ratio ¡ 1 (i.e. *β*-SNR is higher than *α*-SNR) for different segment lengths.

#### 3.3.3. Influence of temporal delays on amplitude envelope cross-frequency correlations

Another result of differential attenuation of separate frequency bands is that correlations between amplitude envelopes across frequencies are influenced by spatial synchronization. An argument for the separation of *α*- and *β*-rhythms into individual components (not stemming from non-sinusoidal waveform) is that only weak amplitude envelope correlations [30] can be found. In simulations, we analyzed amplitude envelope correlations between *α*- and *β*-components, extracted with the corresponding band-pass filtering. Even though the basis functions were generated as a non-sinusoidal waveform with fixed phase delay between the two rhythms, a range of very different correlation values can be observed for individual segments of the compound signal (i.e. a simulation of synthetic EEG data). Fig. 8a shows exemplary time courses for large positive and surprisingly even negative *α*- vs. *β*-correlations. These negative correlations can not be predicted from the amplitude dynamics of individual sources as they only have positive correlations by construction. The observed correlations are dependent on the standard deviation of the mixing coefficient distribution, as illustrated in Fig. 8b, with negative correlations emerging with the increase of the standard deviation of mixing coefficients. The shown examples are for a fixed segment length, but Fig. 8c shows that the effect is present for different lengths of segments.

**Figure 8:**
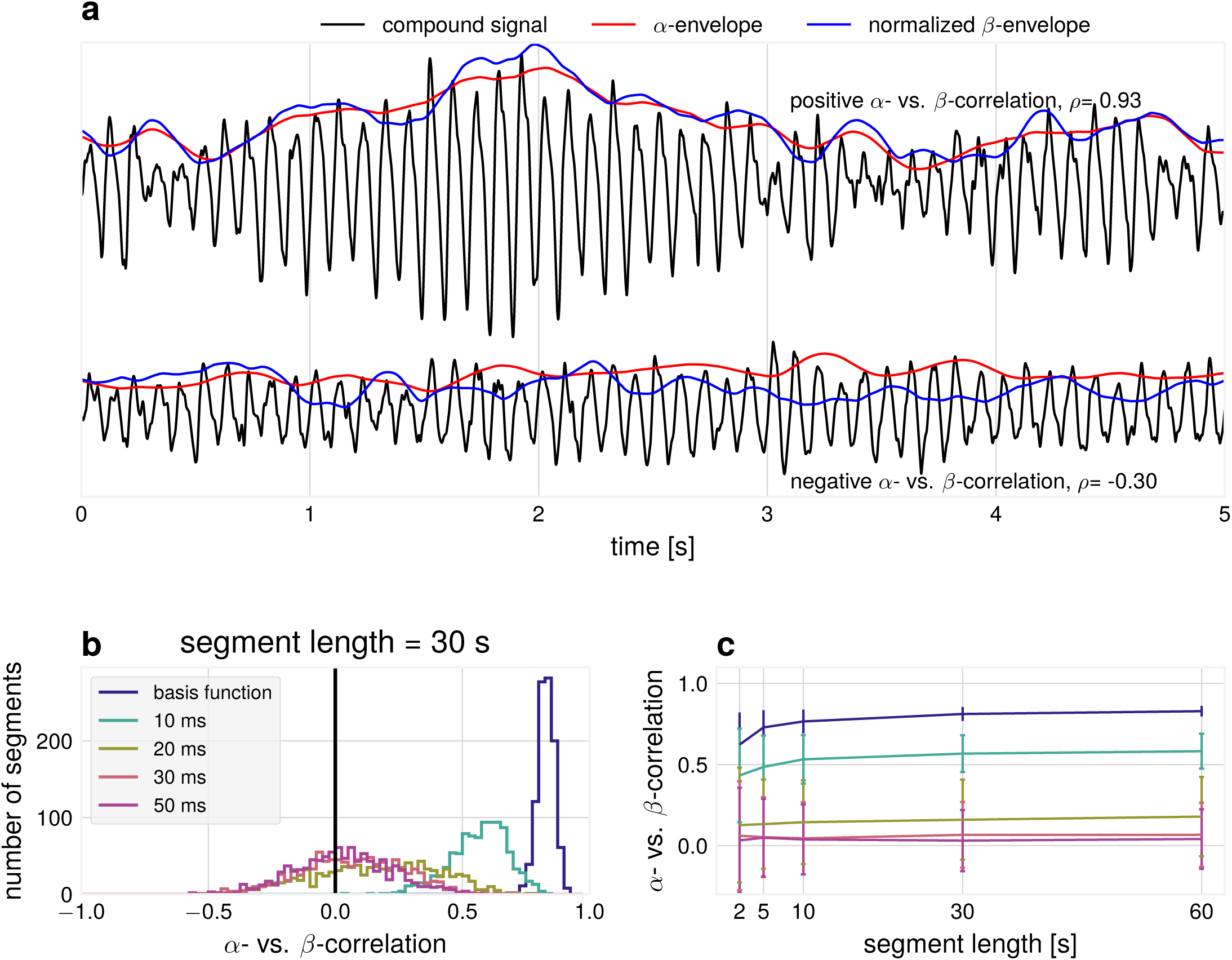
Simulation: relationship between *α*- and *β*-envelopes. (a) Examples for compound signals with positive (top) and negative (bottom) *α*- vs. *β*-envelope correlations. The *β*-envelope was scaled to aid comparisons to the *α*-envelope (*σ* = 10 ms, segment length = 5 s) (b) Spearman rank correlation between synthetic *α*- vs. *β*-envelope time courses over 1000 independent segments. Number of generators = 20. (c) The average correlation as a function of the segment length and standard deviation of the time shift between basis functions.

To validate predictions from simulations, we also quantified *α* vs. *β*-amplitude envelope correlations in empirical data. Fig. 9 shows that amplitude envelope segments as extracted by SSD display larger positive correlations compared to sensor space amplitude envelope correlations. The figure also shows the presence of negative correlations in agreement with the predictions from simulations. Note that with smaller segments one can observe more and stronger negative correlations due to their transient nature.

**Figure 9:**
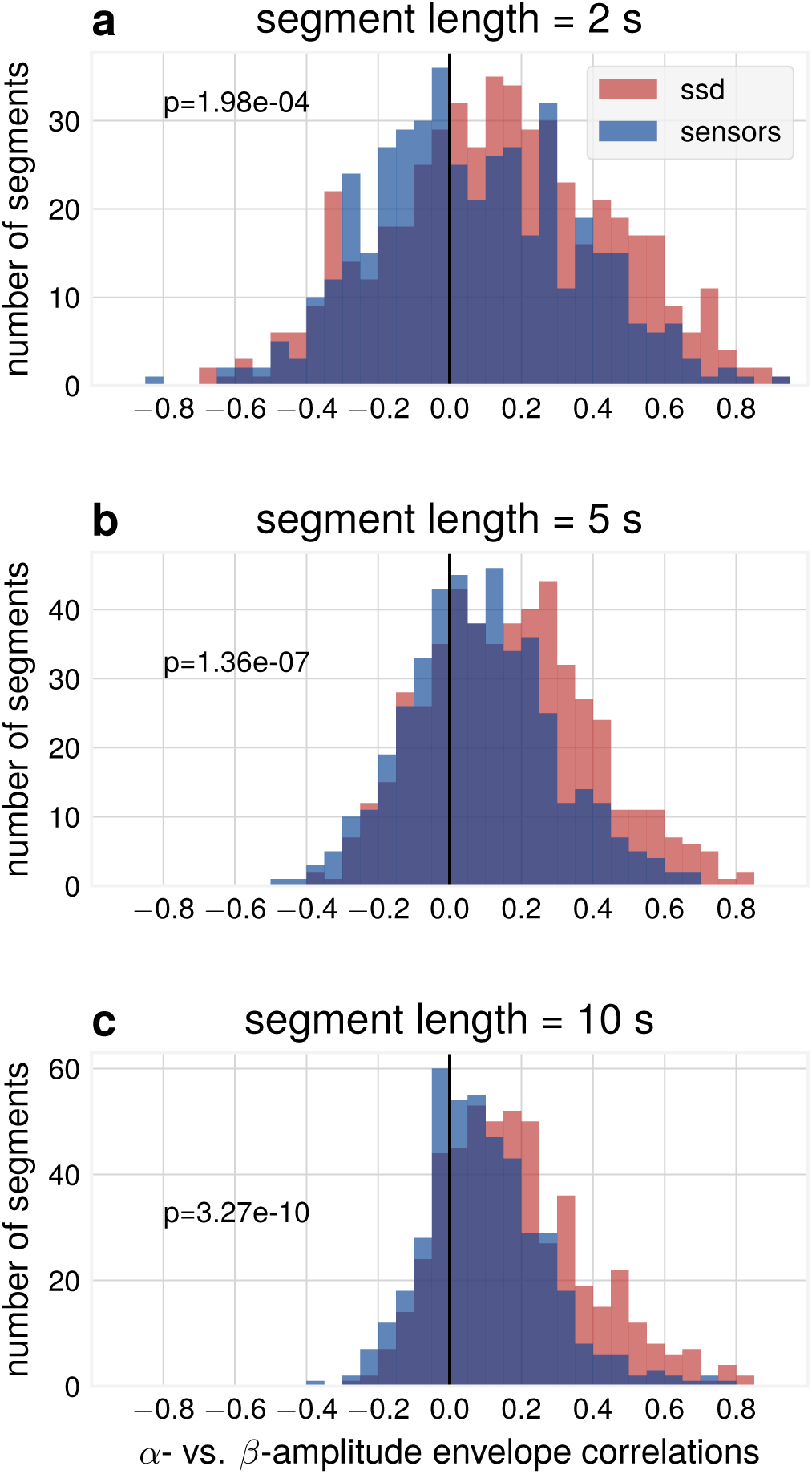
Data analysis: SSD components show increased correlations between *α*- and *β*-envelope time courses compared to sensor space. Subplots are for different segment lengths, (a) 2 s (b) 5 s (c) 10 s. Three sensor space channels (C3, C4, Oz) and three SSD components for each participant, pooled over participants, p-values for Wilcoxon rank sum test (*N*_SSD_=464, *N*_sensors_=374).

## 4. Discussion

In this study we investigated how spatial neuronal synchronization can influence the waveform of neuronal oscillations, affect *α*/*β*-ratios and *α*- vs. *β*-envelope relations. Compound signals become more sinusoidal than their sources for a certain range of temporal delays. We show that the examined measures can be affected solely by these delays even when the basic waveform and spectrum remain the same for the original sources. Moreover, in short segments, *β*-SNR can be larger than the *α*-SNR, through the attenuation of the base frequency. This in turn might relate to the detection in EEG/MEG of *β*-oscillations without the concurrent presence of detectable *α*-oscillations.

Regarding the variability in spectral profiles, different scenarios are possible when estimating *α*-and *β*-relationships arising from the non-sinusoidality of waveforms. Importantly, these diverse spectral profiles can arise from the spatial mixture of non-sinusoidal basis signals with the same waveform. As illustrated in Fig. 10, for the simple scenario with only two sources, different components of a non-sinusoidal waveform can cancel depending on the temporal delay between them. While we primarily focus in this study on the relationships between *α*-and *β*-oscillations, the results can be generalized to the relationships between oscillations at other frequency bands.

**Figure 10:**
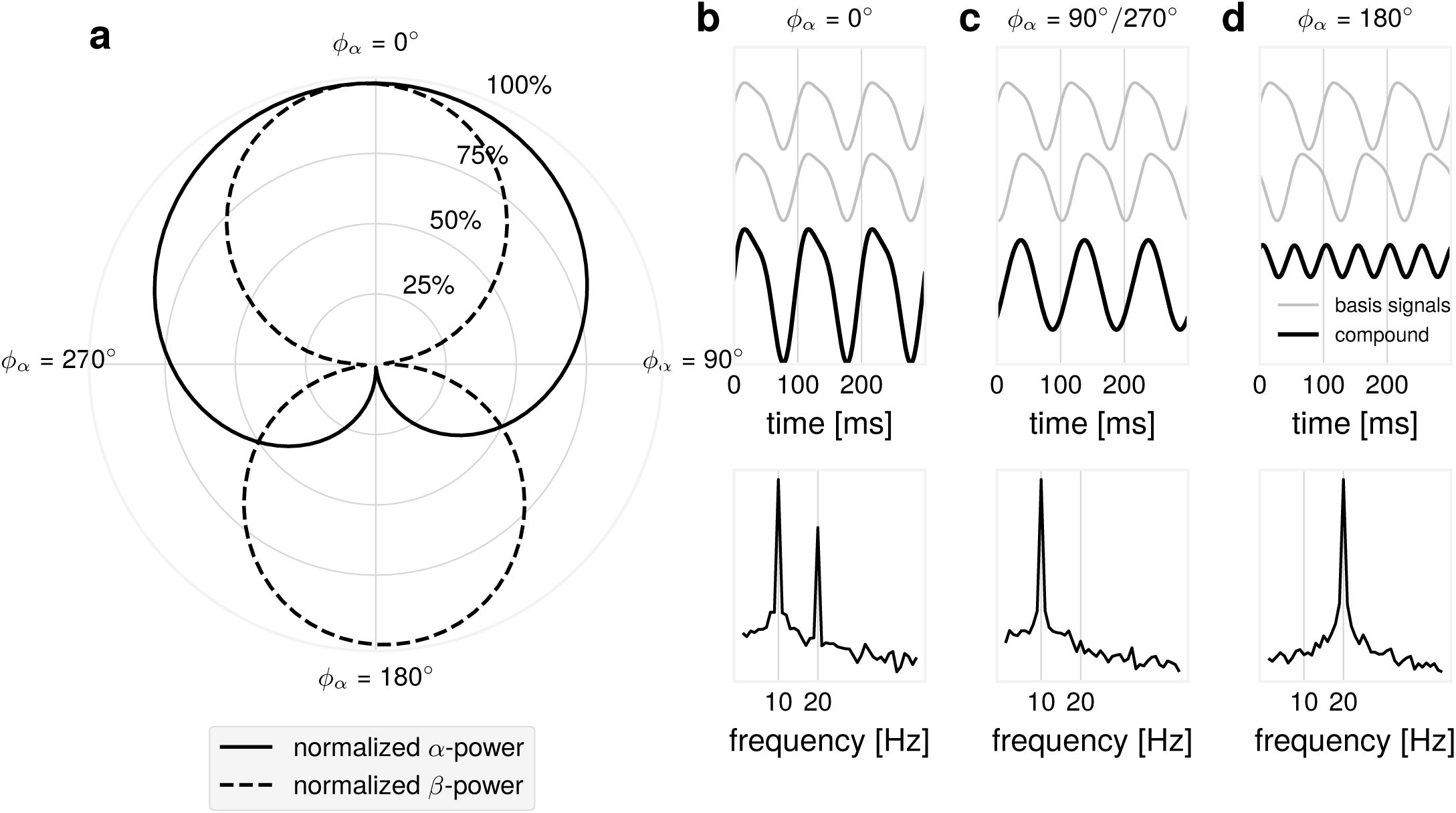
Illustration: possible *α*/*β*-dynamics as function of the time delay for simple synthetic signals. Two non-sinusoidal *μ*-wave signals were mixed with varying time delay *ϕ*_*α*_ between them (*ϕ*_*α*_ = 360^*°*^ is equal to 100 ms, a full cycle of the base *α*-frequency oscillation). A full *α*-cycle corresponds to two full *β*-cycles: *ϕ*_*α*_ = 2 *· ϕ*_*β*_. (a) A polar plot showing *α*-and *β*-power as a function of the time delay *ϕ*_*α*_. *α*-and *β*-power decay differentially as a function of the time delay *ϕ*_*α*_. (b) Time course of the basis signals and the compound signal with the corresponding power spectrum showing maximal *α*-and *β*-power for *ϕ*_*α*_ = 0^*°*^. (c) Time course of the basis signals and the compound signal with the corresponding power spectrum showing attenuation of *β*-power for *ϕ*_*α*_ = 90^*°*^*/*270^*°*^. (d) In the case of *ϕ*_*α*_ = 0^*°*^, in the compound signal, only the *β*-component remains when the *α*-peaks of first basis signal align the the troughs of the second basis signal, causing cancellation.

### 4.1. Limitations

In the present study, we realized the mixing of signals from individual neuronal populations with unitary weights in the simulations. For empirical recordings, data *B* recorded with EEG/MEG can in general be represented as *B* = *L · J*, where *L* is a lead-field matrix and *J* contains dipole currents at different locations. In our simulations, the sources can be assumed to be located close to each other (e.g. < 5 mm) and in practical terms their location and orientation could be considered to be approximately the same thus having the same gain in L matrix. In this way the same gain (unitary or not) for all sources is justified and would lead to similar results. Already on this spatial scale, sources display great dynamical variety [31], with diverse temporal delays [32]. Of course, EEG activity reflects the superposition of a large number of other remote sources, where the mixing of signals at the sensor level would occur with different weights. This, however, would not change one of the main findings of the study qualitatively, namely that the mixing of many non-sinusoidal sources results in more sinusoidal signals.

From our simulations, it follows that if the amplitude weight from one of the sources would be very large (far larger than the weight from other sources), then the signal would remain strongly non-sinusoidal. Only when weights of other multiple sources have sufficient strength and these sources are not synchronized at exactly zero-lag delay [33], only then the superposition of the signals results in more sinusoidal signals. At the level of the remote neuronal populations recorded with EEG, this observation has been confirmed in our study. We showed that ΔCT deviated stronger from 0 for SSD components compared to sensor space data since in the latter case effects of the spatial mixing are more pronounced. Consequently, introducing simulations with different spatial weights would only result in superimposed signals having more non-sinusoidal waveform. Even despite relatively simple but neurophysiologically plausible simulations, we are still capable to show the effects of spatial mixing on waveforms and on complex crossfrequency interactions. While in simulations temporal delays can be specified, these delays are not known for empirical recordings. Therefore, investigating waveform of oscillations can aid in constraining empirical mixing temporal delays.

### 4.2. Implications

Non-sinusoidality in EEG/MEG recordings should be present to a higher degree in signals which demonstrate less spatial mixing. This is the case for instance for many LFP recordings where spatial mixing is restricted to local neuronal populations located in the proximity to the recording electrode [12, 13]. Therefore, non-sinusoidality of the oscillations can be used as a proxy for demixing of neuronal signals recorded with EEG/MEG. Improved methodology will aid in determining functional properties of oscillations with increased sensitivity (not affected by narrow band-pass filtering) when relating oscillatory component to behavioral and stimulation outputs.

Investigating waveform in the temporal domain may aid in an improved determination of phase. A short-coming of current methods for the computation of spatial filters which are based on linear decompositions (SSD, CSP, ICA) is that their solutions are invariant with respective to sign/polarity of the extracted signals. It has been shown that brain states associated with specific phases have differential functional consequences for cortical excitability and plasticity [23, 34, 35]. Therefore, it is important to be able to uniquely define positive and negative peaks of an ongoing rhythm, which is possible when considering measures such the ΔCT. Additionally, the concept of a protophase [26] may aid in describing non-uniform phase velocity and the resulting relationships between cognitive functions and the evolution of oscillations. In fact, as indicated in previous studies [36] duty cycle in neuronal oscillations relates to windows of opportunity for spike transfer between distinct neuronal populations. While 50% duty cycle relates to the same duration of excitatory and inhibitory phases, a deviation from this number (e.g. 30%) can introduce significantly shorter duration of excitatory phase thus providing more precise tuning for the neuronal communication, effectively blocking effects of spikes arriving at the considerably longer inhibitory phase. Spatial mixing in EEG/MEG, leading to more sinusoidal signals, might create an illusion of oscillations with 50% duty cycles while at the source level the duty cycle can be considerably different. When using band-pass filtering non-sinusoidality is removed since only one Fourier component is preserved effectively representing only one frequency and its immediate neighborhood. Behavioral and stimulation effects of such band-pass filtered signal will still be present yet neurophysiological interpretation can be different.

It has been debated whether *α*/*β*-rhythms have a common or separate origin [27, 30, 37]. One of the arguments in favor of both rhythms originating from the same source is that if *α*-and *β*-oscillations are generated by the same neuronal source, producing rhythmic but non-sinusoidal waveform, then one should observe a strong positive amplitude correlation between the two oscillations [30]. This argumentation is based on the linearity of the Fourier transform, as briefly illustrated in the following:

As shown above, our non-sinusoidal signal can be represented as *S* = *α* + *b · β*, with the corresponding Fourier transform of *S* being *F* (*S*). When the amplitude of S is changing in different time segments (multiplied by *A*_*i*_), the corresponding Fourier transform at segment *i*, can be written as: *F* (*A*_*i*_ (*α* + *b · β*)) = *A*_*i*_(*F* (*α*)) + *A*_*i*_ *· b*(*F* (*β*)), which in turns shows that the amplitude of *α*-and *β*-oscillations should covary linearly when the amplitude of S changes by *A*_*i*_. The amplitude of oscillations in different frequency bands can covary for different neuronal sources, but the presence of strong correlations between oscillations at different frequencies with similar spatial topographies is consistent with the idea of them originating from the same neuronal source. Yet, our simulations show that even when a comodulation between *α*-and *β*-oscillations is certainly known to originate from the non-sinusoidal waveform of oscillations, due to the peculiarities of the spatial mixing, it is possible not to observe such positive comodulation. Moreover, surprisingly it is even possible to detect anticorrelation between the amplitudes of *α*-and *β*-oscillations. However, this is entirely due to the effects of spatial mixing of individual signals each of which by itself has only positive correlations between *α*-and *β*-oscillations. Yet, a spatial summation may lead to the occurrence of negative correlations at the sensor level. Importantly, even when using sophisticated spatial filtering techniques such as ICA, SSD, etc. it is unlikely to disentangle such spatial mixing effects originating from the local cortical patches since the resolution of EEG/MEG and even LFP recordings is not sufficient. This also applies to the argument supporting a separate origin of oscillatory components requiring independence of the corresponding temporal dynamics. We have shown that seemingly separate amplitude time courses may not be an indication for the independence of the rhythms, but can also occur when the coupling between different sources changes in the span of only a few hundreds of milliseconds. Whether *β*-events can arise through decoupling of oscillators as in the presented simulations, is a topic for further research. This can reveal insights about mesoscopic brain organization and the interplay of different local rhythms, as extracted by EEG/MEG.

Regarding cross-frequency interactions, our study shows that the amplitude-to-amplitude cross-frequency coupling can also be affected by the non-sinusoidal waveform of the oscillations. For all three types of cross-frequency interactions (phase-to-phase, phase-to-amplitude, amplitude-to-amplitude), spatial synchronization can lead to either very strong or weak indices characterizing cross-frequency interactions, corresponding respectively to a small or rather large jitter in the time delays between neuronal sources (see Fig. 10). This again requires careful interpretation of the obtained data and discussion about the possible effects of spatial synchronization among neuronal populations generating EEG/MEG/LFP signals.

## Acknowledgements

We thank Elena Cesnaite and Keyvan Mahjoory for data preparation.

## Funding

NS acknowledges support through a EXIST Transfer of Research Grant from the German Federal Ministry for Economic Affairs and Energy and from the Quandt foundation. VVN acknowledges support by the HSE Basic Research Program and the Russian Academic Excellence Project “5-100”.

## Declarations of interest

none

